# Structural and molecular characterization of the *Rpv3* locus towards the development of KASP markers for downy mildew resistance in grapevine (*Vitis* spp.)

**DOI:** 10.1101/2021.02.25.432814

**Authors:** Andriele Wairich, Jaiana Malabarba, Vanessa Buffon, Diogo Denardi Porto, Roberto Togawa, Luís F. Revers

## Abstract

*Plasmopara viticola* is the oomycete that causes downy mildew in grapevine. Varying levels of resistance to *P. viticola* across grape cultivars allowed quantitative trait loci to be identified. The *Rpv3* locus is located at chromosome 18, in a region enriched in TIR-NBS-LRR genes, and the phenotype associated is a high hypersensitive response. In this work, we aimed to identify candidate genes associated with resistance to downy mildew on the *Rpv3* locus and to evaluate their transcriptional profiles in a susceptible and a resistant grapevine cultivar after challenging with *P. viticola*. Candidate genes were identified by representational differential analysis and also by functional enrichment tests. Many predicted genes associated with resistance to diseases were found at the *Rpv3* locus. In total, seventeen genes were evaluated by RT-qPCR. Differences in the steady-state expression of these genes were observed between the two cultivars. Four genes were found to be expressed only in Villard Blanc, suggesting their association to the hypersensitivity reaction. Concerning marker assisted-selection for downy mildew resistance, we show the efficient use of a haplotype of SSR markers. Furthermore, based on *Rpv3*-located SNPs between grapevine cultivars contrasting in downy mildew resistance, we developed and tested forty-one new markers for assisted selection. After genotypic and phenotypic evaluations on segregant populations, two markers, Rpv3_15 and Rpv3_33, were considered efficient for downy mildew resistance identification. This study constitutes an in-depth genomic characterization of the *Rpv3* locus, confirms its involvement in resistance against *P. viticola* infection and presents promising biotechnological tools for the selection of young resistant individuals.

## Introduction

Fungal diseases are one of the main constraints for grape production, both quantitatively and qualitatively (Saifert et al. 2018). In humid subtropical areas, as southern Brazil, the most prevalent fungal pathogens responsible for diseases in grapevines aerial parts are *Elsinoë ampelina, Plasmopara viticola*, and *Uncinula necator*, agents that cause, respectively, the anthracnose, downy mildew and powdery mildew diseases (IBRAVIN 2012). *P. viticola* (Berk. Et Curt.) Berl. et de Toni. is a major biotrophic oomycete belonging to the Peronosporaceae (Chromalveolata), known to be a large family of phytopathogenic fungi (Grenville-Briggs and Van West 2005). Grapevine infection by this oomycete starts by the zoospores infiltration via the stomata with the development of a tubular network of hyphae culminating in haustoria proliferation in the leaf mesophyll (Fröbel and Zyprian 2019). After entering the grapevine tissues, it colonizes leaf petioles, shoots, berries and seeds by obtaining nutrients from the infected plant cells with the help of specialized fan-shaped hyphae, that seem to be necessary to overcome physical barriers in plant tissues (Fröbel and Zyprian 2019). The haustoria also allows the exchange of signals involved in the compatibility infection establishment (Polesani et al. 2010). *Vitis vinifera* cultivars are completely susceptible to downy mildew infection, which can cause total production loss by the destruction of inflorescences and/or fruits and by premature defoliation (Erwin and Ribeiro 1996).

In attempting to reduce losses, the intense application of multiple fungicides becomes necessary during each new growth cycle (Cadle-Davidson 2008; Bellin et al. 2009). Even after controlled, *P. viticola* infection may still trigger future crop damages, due to the poor formation of the branches and consequent plant weakened development (Tessmann and Vida 2005; Garrido, L. da R. Sônego 2007). In addition, the disease control measures increase production costs, pose a risk factor for human health and generate environmental impacts in the application areas, such as contamination of soil with chemical residues (Blasi et al. 2011). Besides, there are some *P. viticola* strains with resistance to certain fungicides, which leads to a decrease in the efficiency of disease control (Wang et al. 2013; Blum et al. 2010).

Plant defence responses to pathogens are triggered after the recognition of exogenous chemical signatures by the host which initiate pattern-triggered immunity (PTI). Thereafter, PTI is followed by the immune response signal transduction mechanisms resulting in a significant reprogramming of the plant cell metabolism, which involves changes in gene expression activity (Díez-Navajas et al. 2008; Pinto et al. 2012; Bigeard et al. 2015). This response triggered by plants does not inhibit colonization of the pathogen, but it limits the extent of propagation since the action of disease resistance proteins (R proteins) accelerates and amplifies the innate basal response process. This happens when plants can recognize the pathogen’s effectors via R proteins initiating the effector-triggered immunity (ETI). Plants that are not able to detect these effectors are susceptible the infecting pathogen, resulting in effector-triggered susceptibility (ETS) (Belkhadir et al. 2004; Flor, 1971; Bigeard et al. 2015).

The downy mildew resistance mechanism includes cell wall strengthening, production of antimicrobial compounds such as phytoalexins, the assembly of complex responses such as the hypersensitivity reaction (HR), which is the programmed death of the cells around the infected region, blocking the pathogen progression and the pathogenesis-related (PR) protein synthesis (Polesani et al. 2010). North American grapevine species (*V. riparia, V. cinera, V. labrusca, V. rupestris, V. berlandieri, V. lincecumii* and *Muscadinia rotundifolia*) present variable levels of resistance to *P. viticola* attack (Kortekamp and Zyprian 2003; Unger et al. 2007; Díez-Navajas et al. 2008). More than 200 resistance genes (R genes) were identified in grapevine by the construction of systematic genome maps (Velasco et al. 2007) and the genomic DNA sequencing of *V. amurensis*, an Asian cultivar widely used as an ornamental plant for vertical gardening, and *V. riparia* (Di Gaspero and Cipriani 2003). Many of these genes are located in genomic regions associated with resistance to *P. viticola* in wild *Vitis* species and many have orthologous genes in *Arabidopsis thaliana* that regulate pathways of defence against pathogens (Polesani et al. 2010). Furthermore, the variable levels of resistance to *P. viticola* presented by North American species allowed for the identification of R genes and quantitative trait loci (QTLs) as reported by Merdinoglu et al. (2003), Fischer et al. (2004), Welter et al. (2007), Revers et al. (2010), Blasi et al. (2011), Moreira et al. (2011) and Schwander et al. (2012). Thirty-one QTLs were detected with effect on downy mildew resistance in grapevines, as *Rpv1* (Merdinoglu et al. 2003), *Rpv2* and *Rpv3* (Bellin et al. 2009), *Rpv5* (Marguerit et al. 2009), *Rpv8* (Blasi et al. 2011), *Rpv10* (Schwander et al. 2012), *Rpv11* (Salmaso et al. 2008; Schwander et al. 2012) and *Rpv12* (Venuti et al. 2013). In a more recent report, the discovery of another three loci conferring resistance against *P. viticola* in *V. vinifera* Georgian germplasm, *Rpv29, Rpv30* and *Rpv31*, allowed the identification of potential target genes for grapevine breeding against *P. viticola* based on the possibility to perform crosses with cultivated varieties that present good agronomic traits (Sargolzaei et al. 2020).

The *Rpv3* locus is located in the linkage group (LG) 18 and was identified by different research groups as being associated with a strong HR against downy mildew in resistant individuals (Fischer et al. 2004; Welter et al. 2007; Bellin et al. 2009; Revers et al. 2010; Schwander et al. 2012; Eisenmann et al. 2019; Possamai et al. 2020). This locus accounts for up to 75% of the phenotypic variation, and more than 30 R genes encoding TIR-NBS-LRR proteins and LRR-kinase-type receptor proteins are located within the *Rpv3* region (Velasco et al. 2007; Bellin et al. 2009). The *Rpv3* is also co-localized with the *VviAGL11* locus (AGAMOUS-LIKE 11 (Costantini et al. 2008)), a major QTL for seedlessness, which is a quantitative complex trait (Ocarez et al. 2020).

Currently it is known that the *Rpv3* locus contains at least three different haplotypes (*Rpv3-1, Rpv3-2* and *Rpv3-3*) in segregating populations for downy mildew resistance (Welter et al., 2007; Di Gaspero et al., 2012; Zyprian et al., 2016). The mediated resistance of *Rpv3–1*, which was identified in the cultivar Villard Blanc, is associated with a defence mechanism that triggers synthesis of fungi-toxic stilbenes and HR, resulting in inhibition of pathogen growth and development (Eisenmann et al. 2019). In addition, the transient co-expression of TIR-NB-LRR pair genes, from *Rpv3* locus, in *V. vinifera* leaves activated the HR induced by the pathogen and the sporulation was reduced when compared with leaves from non-transformed plants (Foria et al. 2020).

The TIR-NBS-LRR proteins are related to gene-to-gene resistance response, or the zigzag model (Jones and Dangl 2006). This mechanism is based on the presence of the proteins encoded by the R genes that act in the recognition of specific race effectors, the Avr proteins (Takken et al. 2006; Díez-Navajas et al. 2008; Dry et al. 2010). TIR-NBS-LRR proteins are responsible for the recognition of pathogen-associated molecular patterns (PAMPs), being triggered by effector action/Avr proteins perturbations (Takken et al. 2006). These proteins are composed of three major domains: 1) an amino-terminal variable domain (coiled-coil (CC) or a homologous domain Toll/Interleukin -1-Receptor (TIR), 2) a central nucleotide-binding site (NBS), and 3) leucine-rich repeats (LRR), which is responsible for the recognition of pathogen effectors (Takken et al. 2006). When TIR-NBS-LRR levels increase, they trigger the induction of a rapid defence response that is characterized by calcium and ion fluxes, extracellular oxidative burst, and transcriptional reprogramming in the infected sites and in the peripheral regions, culminating with the HR, ceasing the growth of the pathogen (Belkhadir et al. 2004; Jones and Takemoto 2004; Greenberg and Vinatzer 2003). Therefore, these proteins are determinants of the specificity of the immune response (Takken et al. 2006; Díez-Navajas et al. 2008; Dry et al. 2010).

We chose to work with the Cabernet Sauvignon cultivar because it is one of the most prestigious *V. vinifera* varieties being cultivated in all producing regions because of its capacity to maintain aromas and flavours, independently of the region where it is cultivated (Figueira 2013). However, this variety is sensitive to a series of biotic stresses, for example, to the attack of grapevine pathogens such as *P. viticola* and *U. necator* (Figueira 2013). Contrasting to ‘Cabernet Sauvignon’, the downy mildew resistant cultivar selected for this study, Villard Blanc, also known as Seyve Villard 12,375, is a member of the Seyve Villard family (INRA et al. 2013). ‘Villard Blanc’ is a complex hybrid composed by accesses of six *Vitis* species: *V. aestivalis, V. berlandieri, V. cinera, V. lincecummi, V. rupestris* and *V. vinifera* (INRA et al. 2013). This cultivar presents a strong HR that does not allow the spread of infection beyond the infected cell, which may delay the growth of the pathogen in many interactions, particularly those involving haustoria parasites (Jones and Dangl 2006).

In this work we aimed to identify candidate genes associated with resistance to downy mildew at the *Rpv3* locus in grapevine and evaluate their transcriptional profiles in the susceptible cultivar, Cabernet Sauvignon, and in the resistant cultivar, Villard Blanc, after challenging with *P. viticola*. Moreover, we used SSR markers and KASP ™ genotyping assay to identify single nucleotide polymorphisms (SNPs) or insertions and deletions (INDELs) for the evaluation of the biotechnological applicability of candidate markers, located within the *Rpv3* locus and associated to the resistance phenotype.

### Materials and methods

### Plant material

#### P. viticola *challenge assay prior to gene expression analysis*

Leaves from the cultivar Isabel (*V. labrusca*) naturally infected with downy mildew were harvested from a vineyard in the district of Faria Lemos (29°06’13,09 ‘‘ South and 51°36’26,10 ‘‘ West −362 meters altitude) in Bento Gonçalves – RS, Brazil. This vineyard was subjected to no phytosanitary treatments. The leaves were humidified and stored in a humid chamber at 25 °C and in the absence of light, until sporangia formation. For *P. viticola* challenge assay, 20 young branches of ‘Cabernet Sauvignon’ and 20 young branches of ‘Villard Blanc’ were harvested at Embrapa Uva e Vinho headquarters (29°09’48’’ South and 51°31’53.95” West – 640 meters altitude) and rooted on a Styrofoam box. For optimum *P. viticola* growth, dark and humid incubation chamber was created to ensure high humidity conditions due to the necessity of free water for the production and release of sporangia, formation of zoospores, and subsequent infection of healthy tissues (Tessmann and Vida 2005). Furthermore, to guarantee the assay reproducibility the temperature for fungal growth was adjusted at 25°C for sporangia formation in up to eight hours, following (Ribeiro 2001). A suspension containing 3×10^5^ sporangia mL^-1^ of *P. viticola* was sprayed onto the abaxial surface of leaves. This experiment was carried out in biological triplicates, under controlled environmental conditions (temperature of 25 ± 2 °C and 100% relative humidity). Ten leaves per biological replicate were sampled at 0 (time which was used as a control) 6, 12, 24, 48 and 72 hours after inoculation (HAI). These leaves were immediately frozen in liquid nitrogen and stored at −80 °C until processing.

#### *Evaluation of* P. viticola *resistant genotypes*

Ninety-four grapevine genotypes from a cross between ‘Villard Blanc’ X ‘Crimson Seedless’ (SF-VB population) and segregating for downy mildew resistance were used for SSR markers evaluation. For the Kompetitive allele specific PCR (KASP) markers, two hundred genotypes from a self-fertilized population VB (‘Villard Blanc’ X ‘Villard Blanc’) were used. Moreover, ‘Villard Blanc’ leaves were used for resistance control evaluation and leaves from ‘Cabernet Sauvignon’ were used as a positive control of the infection. For phenotyping their resistance to *P. viticola*, sixteen leaf discs of 1 cm in diameter were detached from each genotype and a suspension containing 3×10^5^ sporangia mL^-1^ of *P. viticola* was sprayed onto the abaxial surface of the discs. Next, they were incubated in Petri dishes with one layer of filter paper under humid conditions, at 25°C and medium light (∼150 µmol m^-2^ s^-1^). The phenotypic classification of downy mildew resistance was determined according to the descriptor OIV - 452 (Anonymous 1983), in which the most susceptible genotypes are classified as 1 and the most resistant, as 9. Genotypes are considered resistant if scored 7 to 9. Disease progression was monitored for nine days. From five days post-inoculation (DPI) onwards, photographs from the dishes were taken daily until the ninth day and a score was given for each genotype by the end of the experiment (9 DPI). These experiments were done in two distinct years.

#### Representational difference analysis (RDA)

Young branches of ‘Villard Blanc’ were challenged with *P. viticola* sporangia or mock solution as described above. ‘Cabernet Sauvignon young branches were used as an infection positive control. ‘Villard Blanc’ leaf samples were obtained at 0, 6, 12, 24, 48 and 72 HAI. Total RNA was isolated (as described in *Plant RNA extraction* section) and equimolar aliquots from each sample point were assembled and used for the synthesis of a cDNA pool. This pool was assayed to obtain a collection of differentially expressed genes using representational difference analysis (RDA) (Hubank and Schatz 1994). Two steps of subtractive hybridization were performed as described in Costenaro-da-Silva et al. (2010). First, an RNA pool from mock-inoculated ‘Villard Blanc’ was used as driver and an RNA pool from downy mildew-inoculated ‘Villard Blanc’ was used as tester. In the second subtraction, the driver was the same and tester was the differential product of the first subtraction. Amplicons from the second subtraction were cloned into pGEM-T Easy vectors (Promega, Madison). A total of 2,229 transformants were obtained, of which 384 were randomly chosen for sequencing. A total of 345 ESTs were found and their sequences were aligned, via BlastN, against the grapevine cDNA genome 12Xv1 (Jaillon et al. 2007). From the 345 EST sequences, 168 corresponded to ribosomal RNA and were excluded from further analysis. The remaining 177 sequences were annotated and their data is publicly available through the GenBank database with accession number JZ984004 and JZ984178.

#### P. viticola *resistance-related candidate genes selection*

For candidate genes selection, Fisher’s test was performed using the software Blast2Go (B2G) (Götz et al. 2008). All genes located at the distal portion of chromosome 18, corresponding to the *Rpv3* locus, were evaluated for enrichment in functional annotation terms (Gene Ontology - GO) related to resistance, taking the entire grapevine gene set as a reference. RT-qPCR direct and reverse primers specific for each candidate gene were designed manually, limiting the size of amplicons to the maximum of 200 base pairs. The candidate genes were named according to their position at the *Rpv3* locus in the 12Xv1 version of the grape genome (Jaillon et al. 2007).

#### Plant RNA extraction

Total RNA was isolated from grapevine leaves by LiCl precipitation using the Zeng and Yang (2002) protocol scaled to 2 mL volumes. Each sample extraction was performed in triplicate and final volumes were pooled before the LiCl precipitation step. Genomic DNA was removed using the TURBO DNA-free Kit (Ambion, Foster City) according to the manufacturer’s protocol. RNA integrity and quantity were monitored by agarose gel electrophoresis and spectrophotometric quantitation, respectively.

#### RT-qPCR analysis

Complementary DNAs were synthesized using the GeneAmp RNA PCR Core Kit (Applied Biosystems, Foster City) according to the manufacturer’s instructions. The gene-specific primers were designed using the OligoAnalyzer 3.1 tool (IDT, http://www.idtdna.com) with the standard settings of 0.2 µM of oligo concentration, 1.5 mM of MgCl_2_ and 0.2 mM of dNTP (Supplementary Table 1). RT-qPCR was performed on a StepOnePlus Real-Time PCR System (Applied Biosystems, Foster City). SYBR Green (Invitrogen, Carlsbad) was used to monitor dsDNA synthesis. Each biological sample was analyzed in technical quadruplicates. Cycling consisted of one step at 95 °C for 10 min followed by 40 cycles of 95 °C for 15 s, 60 °C for 1 min, and finished by a dissociation curve between 60 °C and 95 °C. The specificity of PCR amplification was assessed by the presence of a single peak in melting curves, visualization of single amplification products of the expected size in 3% ethidium bromide-stained agarose gel electrophoresis and sequencing of the amplicons. Primer efficiency was calculated using LinRegPCR software (version 11.0) (Ruijter et al. 2009). Mean relative gene expression was calculated by the (Pfaffl 2001) method employing *ACTIN* (GenBank EC969944) as a reference gene (Reid et al. 2006) and samples harvested at the onset of treatments (0 HAI) were used as a calibrator. All data are represented as averages of three biological replicates and four technical replicates. Statistical analysis was performed using one-way ANOVA followed by Dunnett’s test (p <0.05), due to comparison of treatments with a control, in which the 0 HAI time-point of each cultivar was used as a control for comparison to the following time-points of the same cultivar samples.

#### SSR markers evaluation

Each individual progeny DNA sample from the ‘Villard Blanc’ X ‘Crimson Seedless’ cross was purified as described by Lefort and Douglas (1999) and used in PCR amplifications for each locus as described by Revers et al. (2010). The SSR markers P2_VVAGL11 (Mejía et al. 2011; F-5’ TGTACACCAATACGGGTTTCAT 3’ and R-5’ GTTTGCTGGATTTCCGATGT 3’), P3_VVAGL11 (Mejía et al. 2011; F-5’ CTCCCTTTCCCTCTCCCTCT3’ and R-5’ AAACGCGTATCCCAATGAAG 3’), VMC7F2 (GenBank BV005171; F-5’ AAGAAAGTTTGCAGTTTATGGTG 3’ and R-5’ AGATGACAATAGCGAGAGAGAA 3’) and VVIN16 (GenBank BV140662; F-5’ ACCTCTATAAGATCCTAACCTG 3’ and R-5’ AAGGGAGTGTGACTGATATTTC 3’) UDV108 (F-5’ TGTAGGGTTCCAAAGTTCAGG and R 5’CTTTTTATATGTGGTGGAGC 3’) were used. The amplicons were resolved in 6% polyacrylamide gel and stained with silver nitrate as described by Creste et al. (2001). Deviations between the observed and expected genotype segregations and the possible associations between the phenotypes and the alleles evaluated were tested by concordance and chi-square independence (χ ^2^).

#### SNP markers evaluation

The forty-one SNP-type markers tested in this work were chosen due to their position on the Rpv3 QTL and based on the GrapeReSeq_Illumina_20K chip (https://urgi.versailles.inra.fr/Species/Vitis/GrapeReSeq_Illumina_20K). Leaf discs from the two hundred genotypes from the self-fertilized population VB were shipped to LGC Genomics (http://www.lgcgroup.com, England) to perform the KASP ™ genotyping technique.

#### DNA amplicon sequencing

All the DNA amplicons from RT-qPCR were sequenced and analyzed in an ABI Prism^®^ 310 Genetic Analyser (Applied Biosystems, Foster City) using standard sequencing protocols described in Falavigna et al. (2014). Sequence analysis was carried out with DNA Sequencing Analysis Software v5 (Applied Biosystems, Foster City) and MEGA7 software (http://www.megasoftware.net/home). Sequences were compared to the grapevine reference (‘Pinot Noir’ PN40024) genome 12Xv1.

## Results

### Identification and characterization of associated resistance genes for downy mildew disease

In this study we started by looking for differentially expressed genes after challenging leaves of ‘Villard Blanc’ and ‘Cabernet Sauvignon’ with *P. viticola*. Amongst all specific cDNAs from the RDA approach, two possible candidate genes were identified by their GO terms related to plant-defence response, VIT_04s0008g03400 and VIT_15s0048g02530. The first one is an ethylene-responsive transcription factor from the DREB sub-A-4 family of ERF/AP2 transcription factors (Agudelo-Romero et al. 2015). Our results show that when compared to 0 HAI, this gene had a high decrease in relative expression at 6 and 12 HAI, and kept a low expression in all time-points for both cultivars (Fig.1a). The other candidate, VIT_15s0048g02530 is predicted to be a hairpin-induced 1 gene and its putative orthologous in Arabidopsis is a Late embryogenesis abundant (LEA) hydroxyproline-rich glycoprotein family (AT4G01410), which was described to be involved with defence response against *Pseudomonas syringae* 6 HAI and with an expression peak at 24 HAI (Winter et al. 2007). After challenging with *P. viticola*, VIT_15s0048g02530 presented a significant increase in relative expression at 6 and 12 HAI in both cultivars. Moreover, no differences in expression were observed between cultivars for both genes after 24 HAI (Fig.1a-b). The expression profile patterns of both candidate genes retrieved from the RDA approach were compatible with the strategy envisaged in the experimental design.

Apart from our RDAs results, we further characterized the transcriptional profile of genes described in the literature to be associated with defence response against fungi: Stilbene Synthase (VIT_16s0100g00990), β-1,3-glucanase (PR3, VIT_08s0007g06060) and NDR1/HIN1-like (VIT_16s0098g00890) (Fig.1c-e). The response of these selected genes triggered by the downy mildew challenge assay helped us to point to the expected biological reaction upon fungi infection. The observed expression profile of Stilbene Synthase gene was similar for both cultivars, with an increase in relative expression at 6 HAI, and with significantly higher levels at 12 HAI for ‘Cabernet Sauvignon’ (Fig.1c). For the β-1,3-glucanase gene, the induction of gene expression in ‘Cabernet Sauvignon” started at 12 HAI with *P. viticola*. Nevertheless, this induction was significant in all posterior time-points when compared to 0 HAI, and only for ‘Cabernet Sauvignon’ samples (Fig.1d). For NDR1/HIN1-like gene, the effect of inoculation time was similar between the resistant and susceptible cultivars and the expression peak occurred at 6 HAI, with no significant differences after this time-point (Fig.1e). NDR1/HIN1-like is a probable orthologue of the Arabidopsis NDR1 gene which belongs to the NHL family (for NDR1/HIN1 like) (Dörmann et al. 2000). AtNDR1 is involved in the induction of systemic acquired resistance salicylic acid-dependent, after pathogen infection and many CC-NBS-LRR proteins require NDR1 to trigger multiple signalling defence pathways in Arabidopsis (Century et al. 1997).

**Fig. 1.**
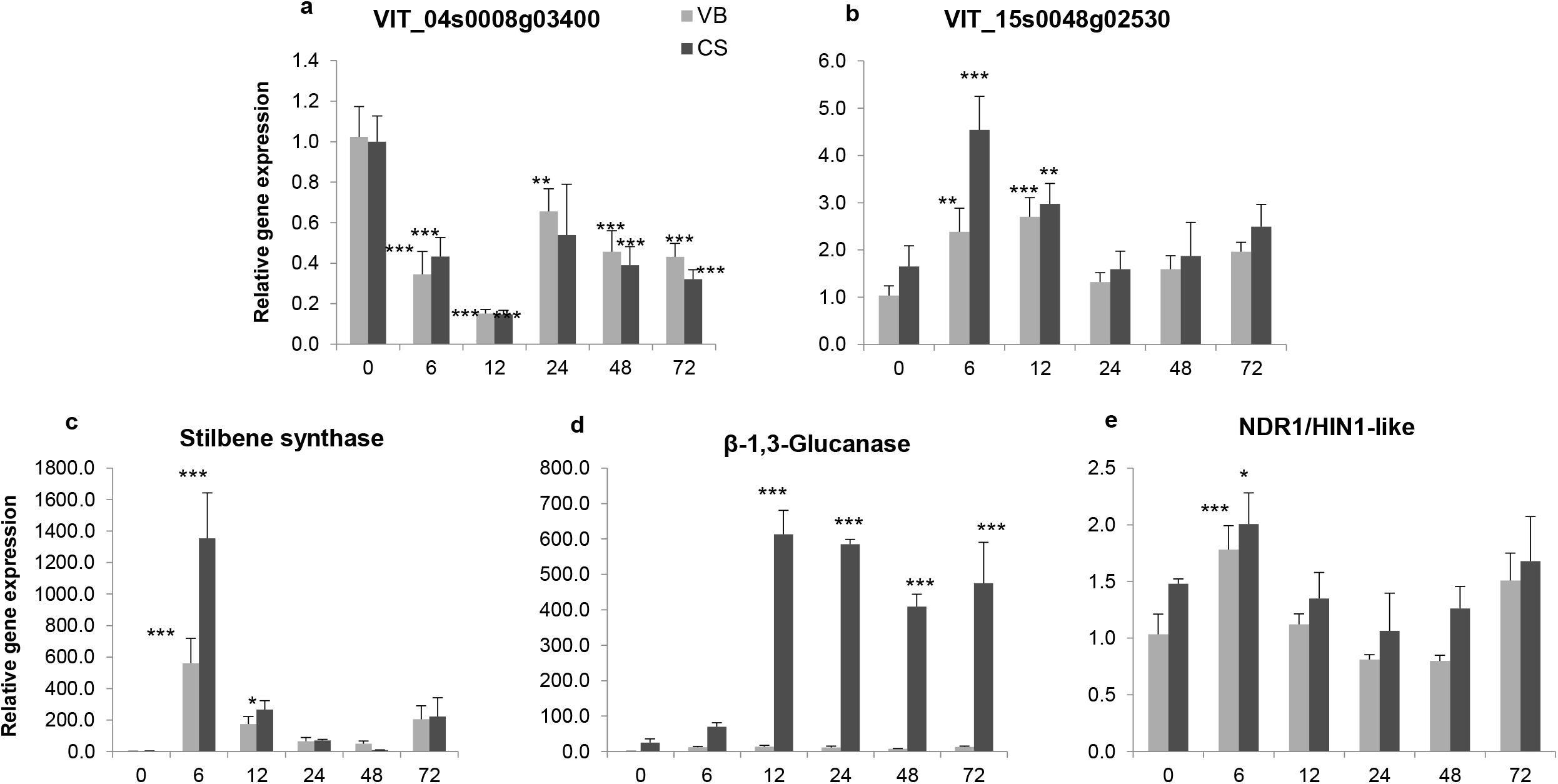
Transcriptional profile of candidate genes after *P. viticola* inoculation. Candidate genes selected from the RDA (a and b). Candidate genes from the literature (c to e). c) Stilbene Synthase (VIT_16s0100g00990), d) β-1,3-glucanase (PR3, VIT_08s0007g06060) and e) NDR1/HIN1-like (VIT_16s0098g00890). Actin gene expression was used as a reference. The X axis shows hours after *P. viticola* inoculation (HAI). VB= Villard Blanc; CS= Cabernet Sauvignon. Standard errors are shown for each time point. One-way ANOVA followed by Dunett’s test, in which the 0h time-point of each cultivar was used as a control for comparison to the following time-points, * p< 0,05; ** p< 0,001; *** p<0,0001.

### *Identification of resistance genes on* Rpv3 *locus associated with downy mildew defence response*

For better exploring the *Rpv3* locus (LG18) we aimed to identify and test possible candidate genes for downy mildew resistance. By performing a Fisher’s test, we observed that the genomic region located in the *Rpv3* locus, based on the reference genome PN40024,is enriched with genes associated with defence responses, programmed cell death, signal transduction, immune system processes and responses to stress (Supplementary Table 2). About 70 genes were found at the *Rpv3* locus, of which 40 had at least one of the TIR-NBS-LRR domains, while eight genes had all three domains. From this set of 40 predicted candidate genes, 23 were selected for RT-qPCR evaluation (Supplementary Table 2), 8 of them for having the 3 TIR-NBS-LRR domains and 15 for having more GO terms associated with programmed cell death and defence responses, besides being located close to the seedlessness selection molecular marker VMC7F2 (Cabezas et al. 2006) and having at least one of the TIR-NBS-LRR domains. The location of the 23 genes on the *Rpv3* locus selected for the accomplishment of the present work is demonstrated in Fig.2.

**Fig. 2.**
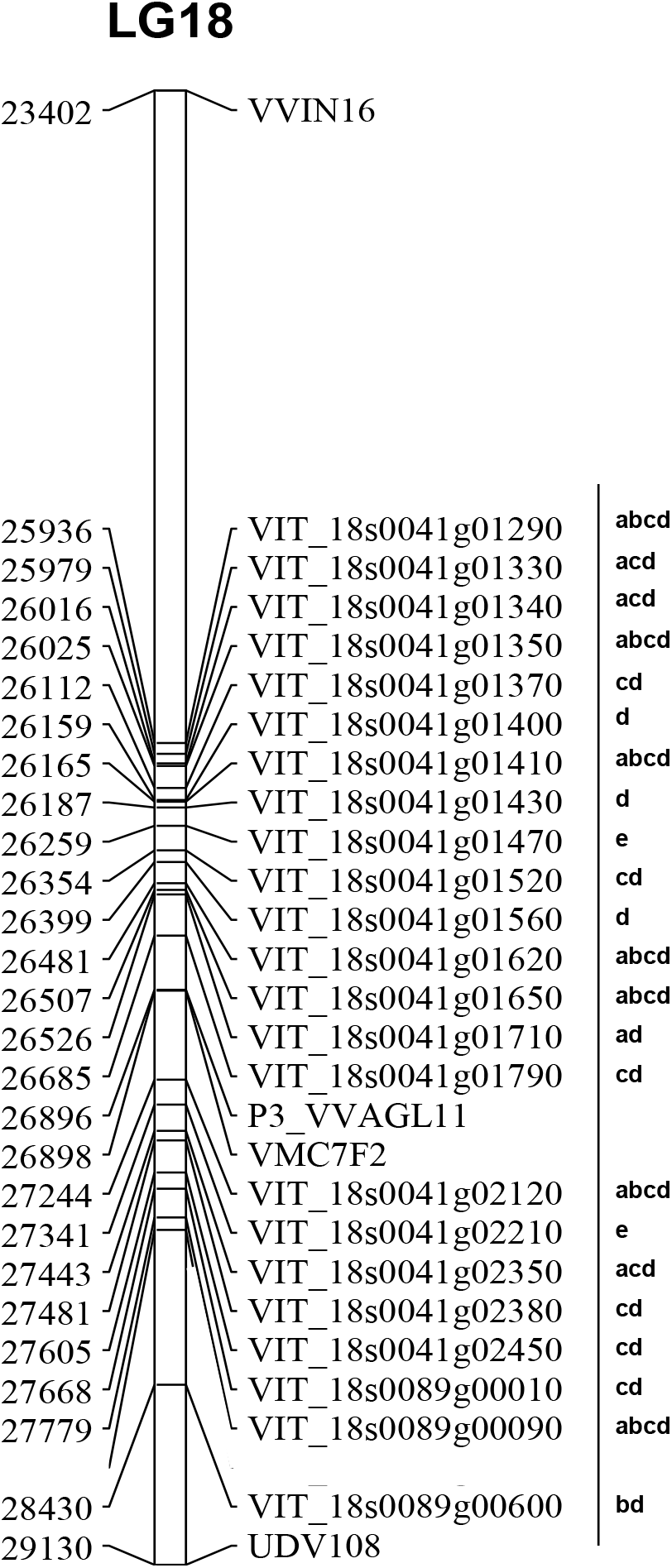
The position of the candidate genes at *Rpv3* locus on the distal portion of chromosome 18. Letters represent the presence of amplification products for each cultivar, Villard Blanc (VB) and Cabernet Sauvignon (CS), by PCR and by RT-qPCR (a= RNA VB; b= RNA CS; c= DNA VB, d= DNA CS and e= not detected/amplified). Numbers on the left are megabases (MB) and chromosome coordinates are from reference genome PN40024.

### *Characterization of* Rpv3 *candidate genes*

To characterize the candidate genes, conventional PCR reactions based on genomic DNA were performed to verify their presence in the genome of both cultivars, as well as to confirm the size of the amplification product of each gene, based on the grapevine genome from cultivar Pinot Noir (PN40024). Of the 23 candidate selected genes, 21 yielded the predicted product after amplification of ‘Cabernet Sauvignon’ DNA samples. On the other hand, ‘Villard Blanc’ was positive for only 16 candidate genes under the conditions tested (Supplementary Fig.1). This allowed us to confirm the presence of most candidate genes in the genomes of both cultivars. A few genes presented differences in the amplification product sizes between cultivars. The gene loci, specific primers and expected amplification products are described in Supplementary Table 1.

### Rpv3 *candidate genes expression in response to P. viticola infection*

Transcriptional profiles of candidate genes were obtained by RT-qPCR after spraying *P. viticola* sporangia on leaves from the susceptible cultivar Cabernet Sauvignon and the resistant one Villard Blanc. *Actin* gene expression was used to normalize the data from all genes and samples harvested at the onset of treatments (0 HAI) were used as a calibrator. Out of the twenty-one pairs of specific primers used to evaluate the expression of candidate genes, nine did not show RT-qPCR amplification. For twelve genes it was observed amplification products of expected sizes. The genetic specificity of the twelve sequenced PCR products was confirmed by sequencing.

The twelve genes that presented expression were divided into three groups, according to their expression pattern: 1) the ones that presented expression in both cultivars samples (seven genes – Fig.3a-g); 2) the ones that only had expression in the Villard Blanc cultivar samples (four genes – Fig.3h-k) and 3) one gene that was only expressed in ‘Cabernet Sauvignon’ samples (Fig.3l).

From the first group, the transcriptional profile of the genes VIT_18s0041g01350, VIT_18s0041g01620 and VIT_18s0089g00090 (Fig.3b, 3d and 3g) demonstrated an increase in relative gene expression after challenge with *P. viticola* in both cultivars. The gene VIT_18s0041g01350 presented a transcriptional profile with high levels for ‘Villard Blanc’ samples with statistical differences after 6 HAI (Fig.3b). Furthermore, for the Cabernet Sauvignon cultivar, the gene VIT_18s0089g000090 was approximately 2X more expressed after 6 HAI (Fig.3g). VIT_18s0041g01620 presented a significant difference in 48 and 72 HAI when compared to 0 HAI showing an approximately 2.5X increase in expression in ‘Cabernet Sauvignon’ (Fig.3d). VIT_18s0041g02120 presented a higher level of transcripts for ‘Cabernet Sauvignon’ samples in most time-points while for Villard Blanc cultivar its expression decreased after *P. viticola* inoculation (Fig.3f). This tendency of differential expression with a decreasing curve happened for other genes, such as VIT_18s0041g01290, VIT_18s0041g01410 and VIT_18s0041g01650 (Fig.3a, 3c and 3e, respectively) for both cultivars.

Interestingly, four genes presented expression only for the resistant cultivar Villard Blanc after *P. viticola* challenge, representing the second group (Fig.3h-k). Genes VIT_18s0041g01330 and VIT_18s0041g01340 showed a gradual decrease in relative gene expression in the period between 0 and 48 HAI with *P. viticola* (Fig.3h-i). Moreover, gene VIT_18s0041g01710 presented a statistical difference in relation to 0 HAI with a significant decrease at all time-points for resistant cultivar Villard Blanc (Fig.3j), while gene VIT_18s0041g02350 presented a variable expression during the experiment time points (Fig.3k).

The VIT_18s0089g00600 gene, that composes the third group, was the only one expressed uniquely in the susceptible cultivar. This gene showed a slight increase in expression after challenge with *P. viticola*, demonstrating a 2.5X higher expression at 72 HAI when compared to 0 HAI (Fig.3l).

**Fig. 3.**
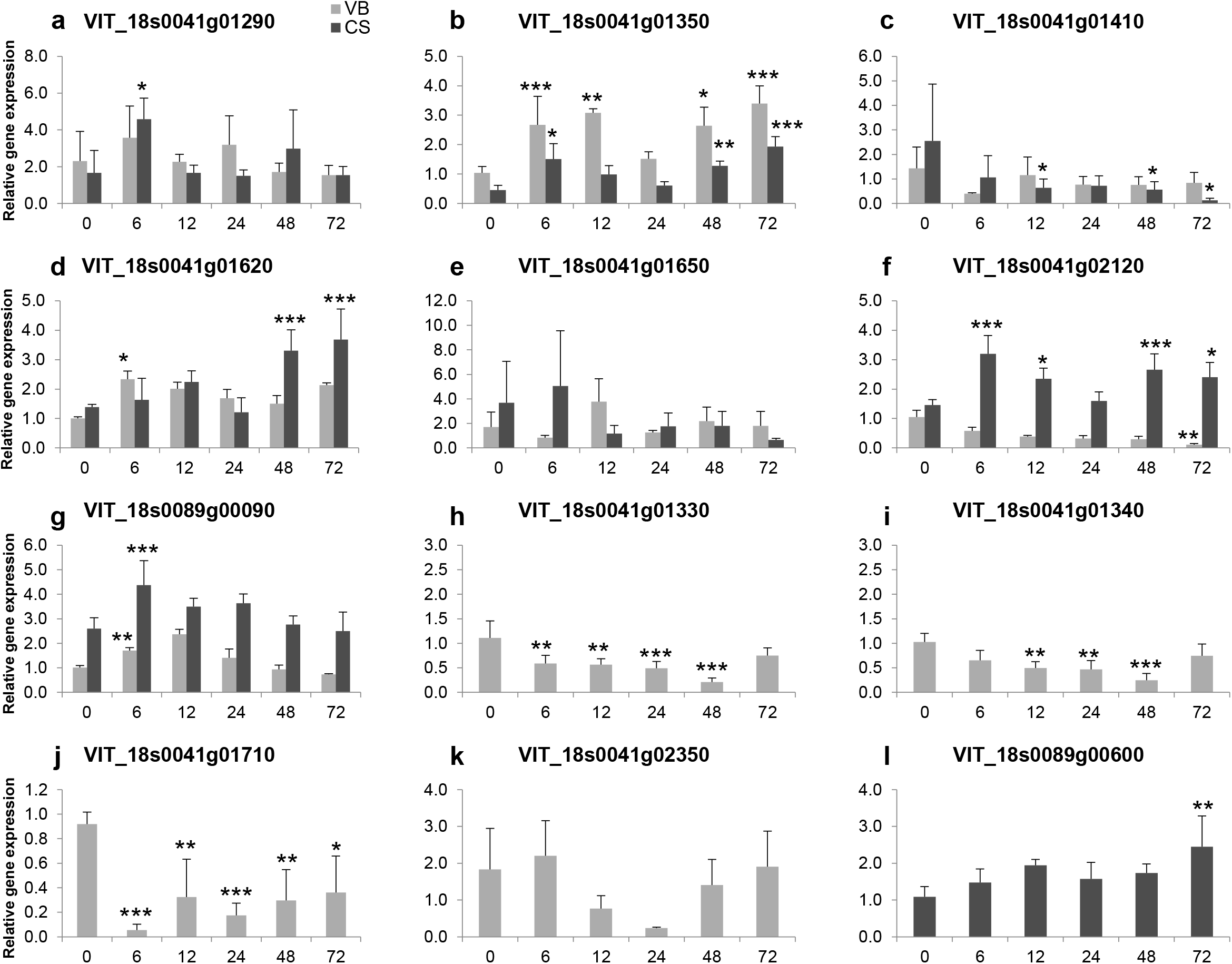
Transcriptional profile of candidate genes at the *Rpv3* locus after *P. viticola* inoculation. Actin gene expression was used as a reference. Axis *x* shows hours after *P. viticola* inoculation (HAI). VB= Villard Blanc; CS= Cabernet Sauvignon. Standard errors are shown for each time point. One-way ANOVA followed by Dunett’s test, in which the 0h time-point of each cultivar was used as a control for comparison to the following time-points, * p< 0,05; ** p< 0,001; *** p<0,0001.

### Evaluation of SSR markers for downy mildew resistance

The co-localization of the *Rpv3* locus and the *SdI* locus (Seed development Inhibitor) gave us the opportunity to test molecular markers that were already established for seedlessness assisted selection. These markers (UDV108, VMC7F2, VVIN16, P2_VVAGL11 and P3_VVAGL11) are known for their efficiency (>80% accuracy) as tools for assisted selection of apirenic grapevines (Doligez et al. 2002; Cabezas et al. 2006; Mejía et al. 2011; Ocarez et al. 2020). For this analysis, we performed the phenotypic characterization of resistant genotypes on ‘Villard Blanc’ X ‘Villard Blanc’ population after a *P. viticola* challenge assay (see Materials and Methods - Fig.4). On our bioassays, we were able to identify reddish-brown tissue of necrotic spots, corresponding to the oil stain, in the upper region of the infected leaves in all the performed assays. Depending on the severity of lesions, plants were scored for downy mildew resistance or susceptibility, with the resistant plants presenting a score between 5 to 9, while susceptible plants present a score of 1 to 3 (OIV - 452) (Fig.4). After performing the χ2 adjustment tests, UDV108 and P2_VVAGL11 did not present the expected segregation and were excluded from the analysis. For the loci VMC7F2, VVIN16 and P3_VVAGL11 four segregating alleles were observed in the genotyped population. Analysis of the phenotypic distribution of the evaluated characters versus the allele frequency of the genotyped SSR markers shows the association between the alleles P3_VVAGL11-185 bp (χ2 calc = 28.9), VVIN 16-154 bp (χ2 calc = 26.81) and VMC7F2-210 bp (χ2 calc = 36.93) with downy mildew resistance (Fig.5). By the evaluation of these marker genotypes, haplotypes were identified for resistance to downy mildew. The use of the haplotype that combines the three SSR markers, VMC7F2, VVIN 16 and P3_VVAGL11, presents 100% accuracy in the selection of resistant individuals in the samples of plants tested, also eliminating potential false positives.

**Fig. 4.**
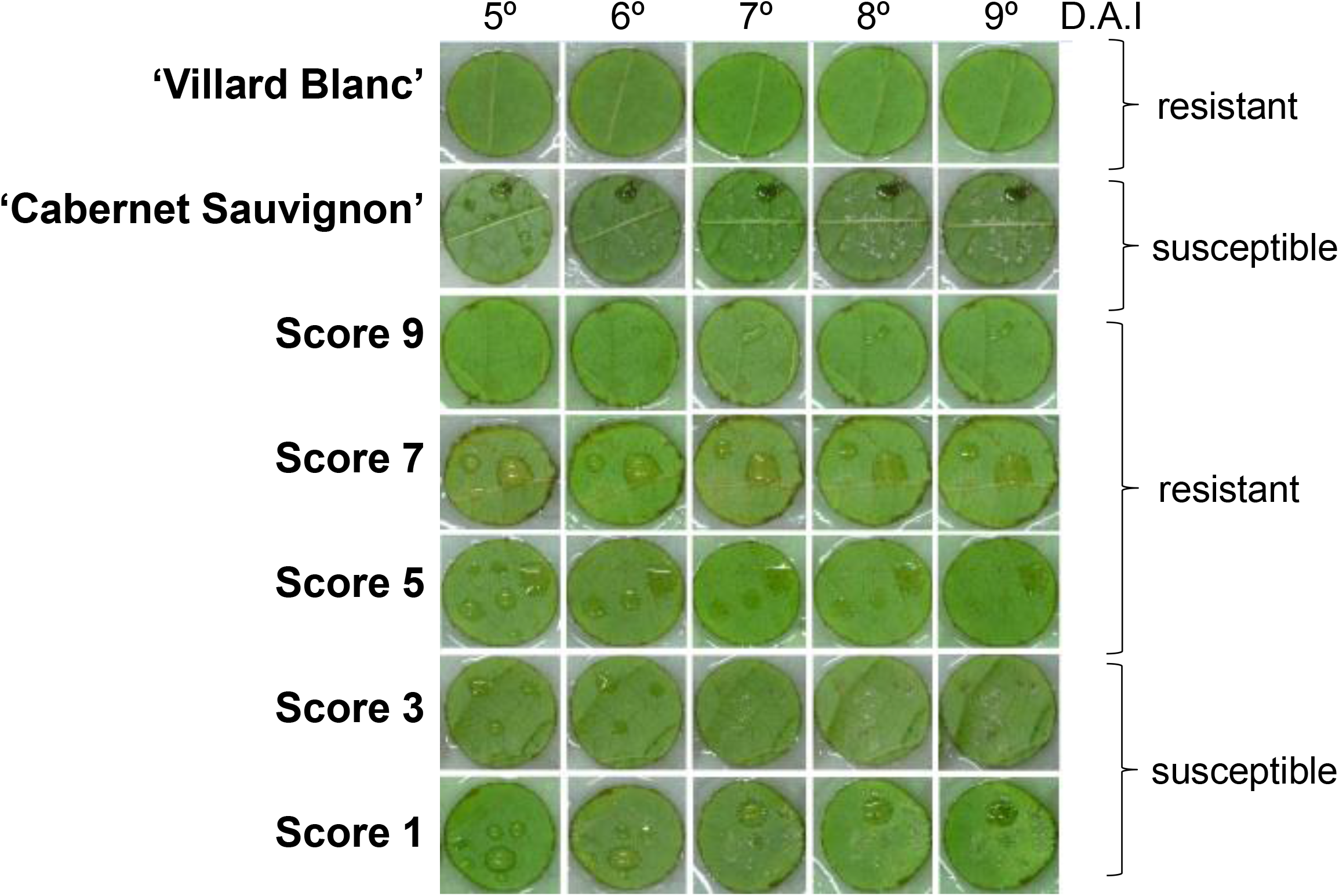
Downy mildew infection progression on leaf discs. Leaf discs were monitored for 9 days after *P. viticola* inoculation. OIV-452 score was used as reference, in which scores 1 to 3 are defined as susceptible to *P. viticola* and scores ranging from 5 to 9 are defined as resistant to *P. viticola*. Leaves from 5 DAI to 9 DAI are represented for selecting genotypes of each score. Leaf discs from the paren ‘Villard Blanc’ were used as the resistance control and ‘Cabernet Sauvignon’ was used as positive control for the infection.

**Fig. 5.**
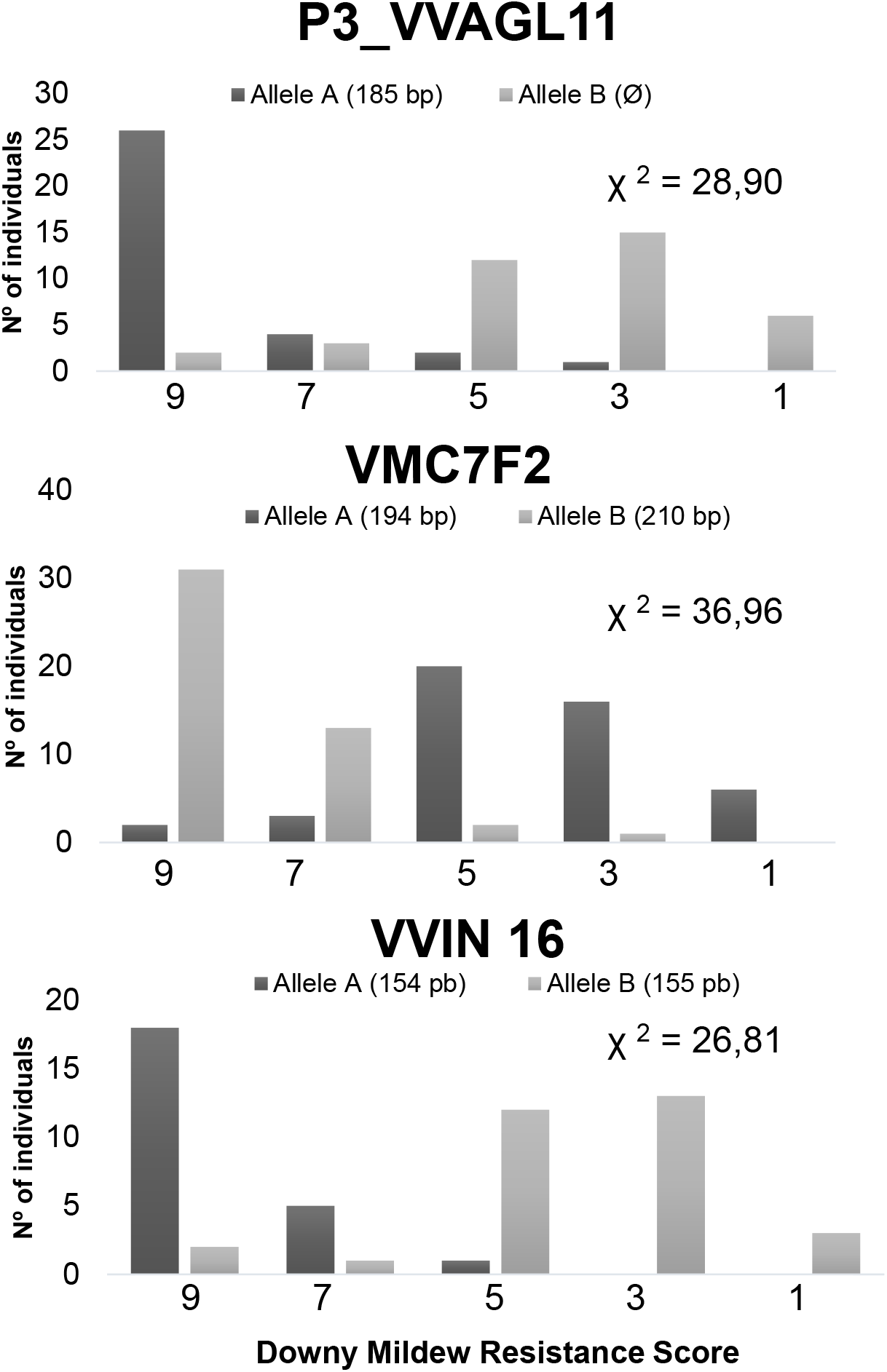
SSR markers evaluated for downy mildew resistance. Number of F1 individuals from population ‘Villard Blanc’ X Crimson Seedless, and their genotypes versus the score for *P. viticola* resistance (OIV - 452).

### *Development and evaluation of SNP markers for assisted selection of resistance to* P. viticola

In order to develop new efficient molecular markers for assisted selection of downy mildew resistance, we chose SNPs that are located within the *Rpv3* locus. These selected mutations were based on a high throughput genotyping of different grapevine backgrounds (Supplementary Table 3). We developed forty-one markers to be tested by competitive allele PCR (KASP ™). Of those, Rpv3_15 and Rpv3_33 loci presented segregating polymorphisms in the genotyped population (VBXVB). The other markers tested were homozygous in this population. Analysis of the phenotypic distribution versus allele frequency clearly showed (p <0.0001) the association between the heterozygous form of the Rpv3_15 (χ^2^ calc= 59.81, position chr18_26844557) and the Rpv3_33 (χ^2^ calc= 119.9, position chr18_27469511) with downy mildew resistance (Fig.6). All the candidate markers tested are represented at chromosome 18 in Fig.7.

**Fig. 6.**
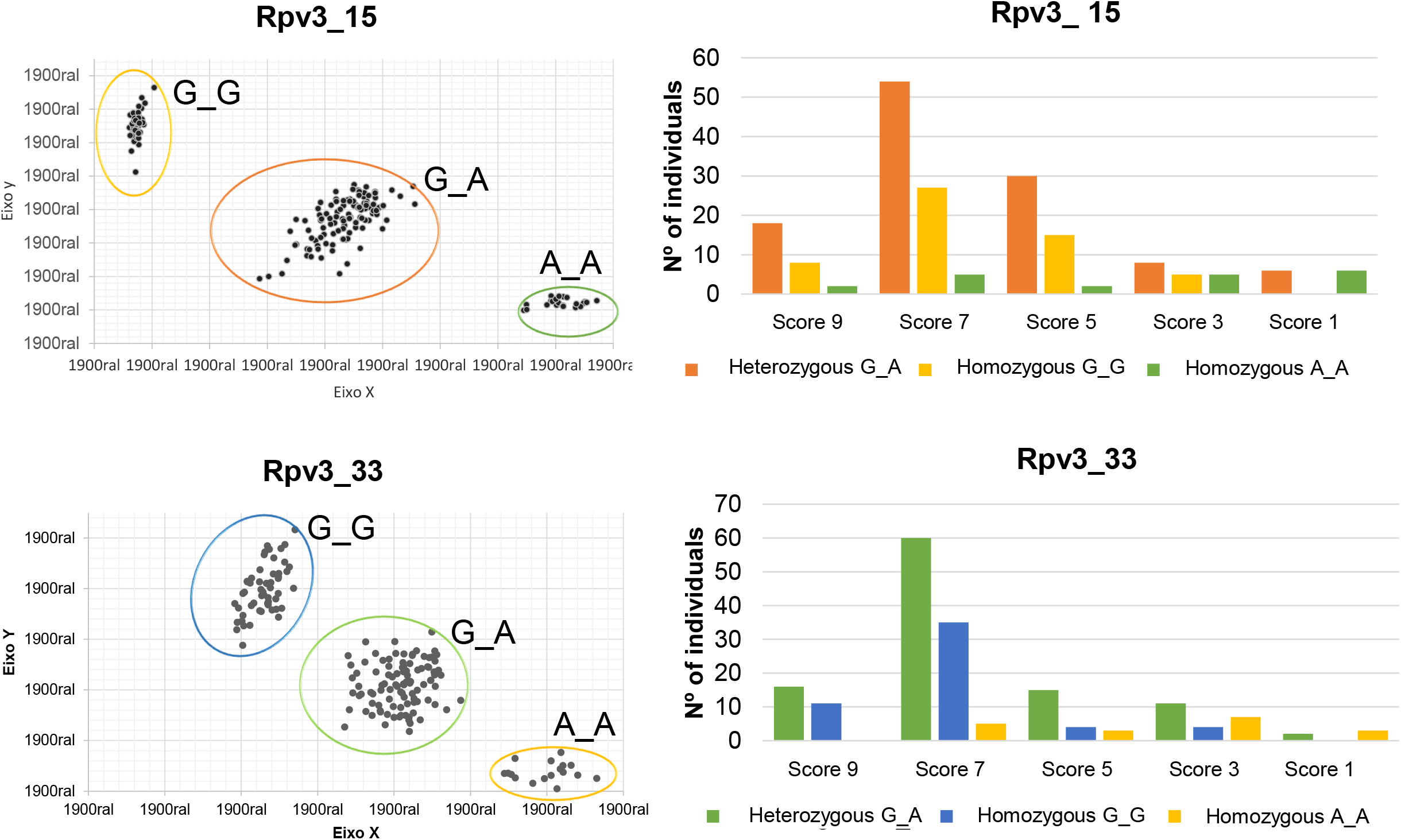
Evaluation of SNP-type markers in downy mildew resistant segregating population. KASP markers depicted on the left panel shows the fluorescence results for FAM and HEX in the axis *x* and *y*, respectively. Right panel demonstrates the relation between the number of individuals and their genotype versus the score for downy mildew resistance (OIV - 452).

**Fig. 7.**
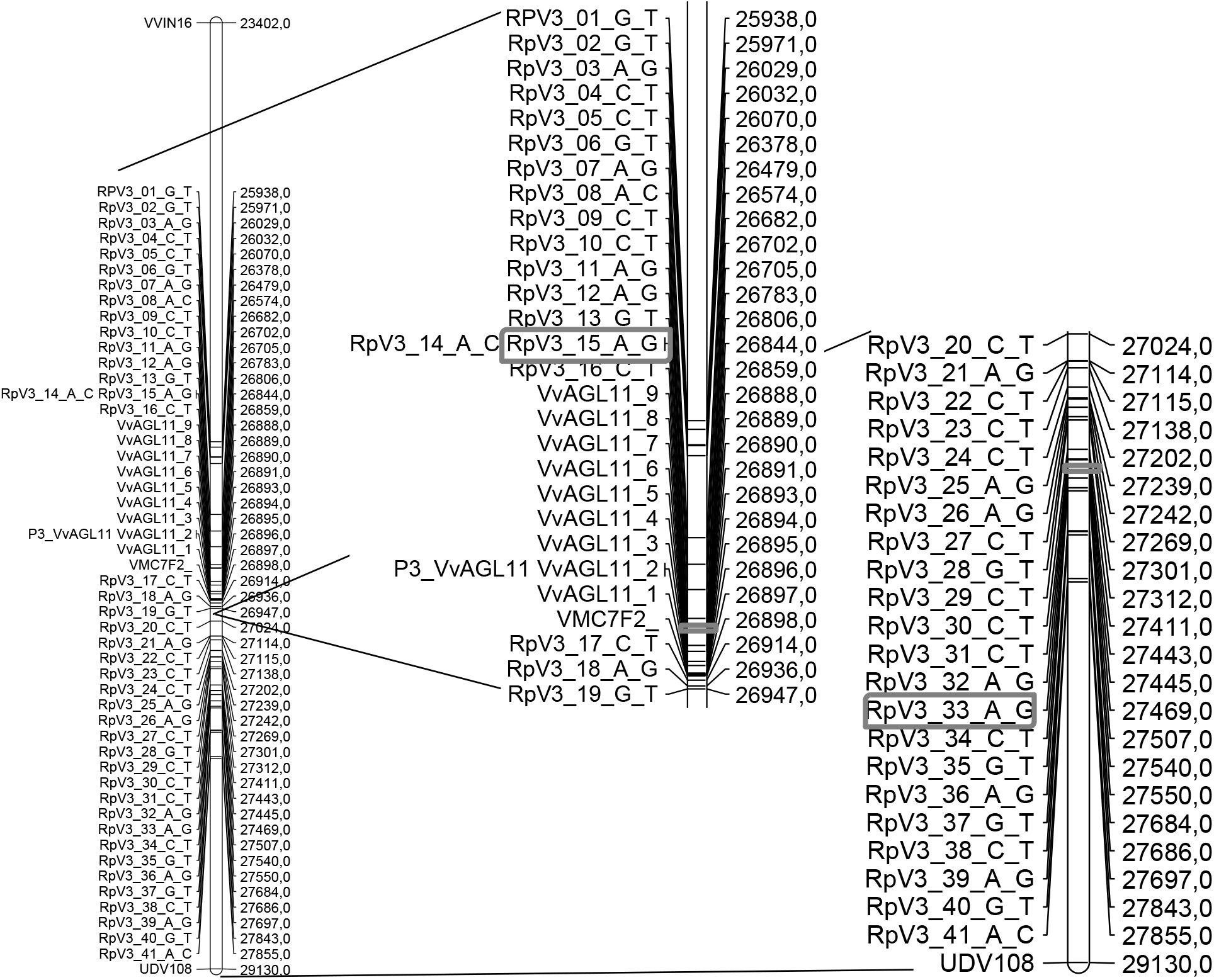
SNP-type candidate marker’s position at the *Rpv3* locus. The markers were developed for competitive allele-specific PCR (KASP) for the evaluation of downy mildew resistant segregation. The SSR markers on the distal portion of chromosome 18 (reference genome PN40024) are the ones tested in this work (VVIN16, P3_VVAGL11, VMC7F2 and UDV108). Markers surrounded by a grey box presented segregating polymorphisms related to downy mildew resistance in the genotyped population.

## Discussion

In the current moment of agricultural modernity it is urgent to understand the molecular and physiological mechanisms of plant defence against a large range of abiotic stresses. And more imperative is then to use this knowledge for the development of sustainable and earth-preserving field practices. In this work we aimed to follow two grape cultivars’ response to *P. viticola* infection using the expression of putative homologs of defence genes as indicators of their relation to downy mildew resistance. During our bioassays that challenged plant leaves or plant leaf discs with *P. viticola* we followed the presence or absence of reddish-brown stains on the upper part of the leaves as a resistance or susceptibility indicator, respectively. These stains are due to the pathogen recognition system which triggers localized resistance reactions such as the hypersensitivity reaction (Durrant and Dong 2004). When in nature, over time, these stains increase in size and unite occupying much of the leaf blade with the tissue in the spot’s area becoming dry, causing the affected leaves to fall prematurely and depriving the plant to generate photoassimilates (Ribeiro 2001; Garrido, L. da R. Sônego 2007).

Our candidate gene expression analysis results demonstrate the intricacy of the downy mildew resistance mechanism. Studies show that the time and intensity with which the infected organism develops the defence process presents a major genotypic component, thus the process of resistance induction is functionally, spatially and temporally complex (Bellin et al. 2009; Pinto et al. 2012). One of the genes selected by the RDA approach, VIT_15s0048g02530, shows that ‘Cabernet Sauvignon’ and ‘Villard Blanc’ had a fast, but transient response against *P. viticola*. In addition, the expression analysis of genes selected from the literature demonstrates the triggering of ‘Cabernet Sauvignon’ defence response. Especially for β-1,3-glucanase transcriptional profile, ‘Cabernet Sauvignon’ presented extremely high levels when compared to ‘Villard Blanc’ showing that some defence pathways are being activated, but their products are not sufficient to prevent downy mildew infection.

We also identified new candidate genes located at the *Rpv3* locus by their predicted function as related to fungi resistance, which is determinant of plant immune response specificity. Not all the performed PCRs showed amplification for both cultivars analyzed (SFig.1). The absence of amplification products for the Villard Blanc cultivar may be related to the fact that this cultivar is a complex hybrid and distantly related to ‘Cabernet Sauvignon’, as a consequence has differences when compared with the inbreeding line PN40024, from which the full genome was sequenced, and from which ‘Cabernet Sauvignon’ is closely related. Future sequencing of the ‘Villard Blanc’ genome could answer this question.

The results comprising genes that were expressed in both cultivars show interesting data for VIT_18s0041g01350, VIT_18s0041g01620 and VIT_18s0089g00090. These transcripts were expressed with significant differences between time-points. According to Blast2GO functional terms mapping, the VIT_18s0041g01350 and VIT_18s0041g01620 are related to processes of apoptosis, ATP binding, innate immune response, transmembrane activity receptor and signal transduction. The gene VIT_18s0089g00090 presents GO terms related to apoptosis processes, transmembrane activity receptors, binding nucleotides and signal transduction. In addition, both VIT_18s0041g01620 and VIT_18s0089g00090 predicted proteins present the three TIR-NBS-LRR domains. The results suggest that these three genes may be related to the effector-triggered immune response process that results in the assembly of the HR in resistant individuals.

A transcriptional profile for the genes VIT_18s0041g01330 and VIT_18s0041g01340 was only visible in the resistant cultivar. Both genes present the three domains, TIR, NBS and LRR, in their predicted proteins, and both were annotated with GO terms related to processes of apoptosis, binding proteins, innate immune response and stress-causing factors, among other processes. Their transcriptional profiles suggest that these genes may be related to the process of immune response triggered by the effector that results in HR. The decrease in gene expression levels, as compared to 0 HAI, presented by these two genes might be related to a part of the infection response that we still do not understand, but this decrease has already been reported in a study by Soanes and Talbot (2008). Moreover, genes VIT_18s0041g02350 and VIT_18s0041g01710 presented a transcriptional profile only in the downy mildew resistant Villard Blanc cultivar after challenge with the pathogen. Both genes present GO terms associated with apoptosis, ATP binding, defence responses and stress, among others processes. Their transcriptional profile suggests that they may be associated with the defence response against *P. viticola*. This set of genes, only expressed in ‘Villard Blanc’, could be responsible for the contribution of the *Rpv3* locus to ‘Villard Blanc’ effective defence response upon downy mildew infection. Our results deviate from the work of Polesani and collaborators (2010) in which they observed that the resistant genotype, in their case cultivar Gloire (*V. riparia*), starts assembling the defence response mechanism against downy mildew in the first 24 hours after contact with the pathogen. This is the period in which the plant innate immune system is activated by a large number of surveillance-type receptors that play a role in detecting the presence of pathogens and transmitting the invasion signal, which is followed by the establishment of the compatible plant-pathogen interaction (Polesani et al. 2010; Andolfo and Ercolano 2015). In this scenario, the structural defence mechanisms of the host has influence at the early interaction between the pathogen and its host, as observed in a multi-year study for the evaluation of resistance of Georgian grapevine germplasm and *P. viticola* (Toffolatti et al. 2016).

In our evaluation of candidate genes transcriptional profiles, it was possible to observe that ‘Villard Blanc’ presented almost a constant expression of the genes tested, except for VIT_18s0041g00090, that 6 HAI had a significant induction. This suggests that the resistant cultivar could be, in a way, always producing the necessary defence response proteins against the pathogen. In the case of *V. vinifera* cultivars, it is known that they are susceptible to the attack of *P. viticola*, although they present defence responses against other pathogens, thus indicating that the defence components specifically against *P. viticola* are not activated in a necessarily short period after infection (Kortekamp 2006). From these data we can hypothesize that the presence of the selected candidate genes in the genome of both cultivars, still demonstrates the absence of one or more components in *V. vinifera* that would be necessary to trigger *P. viticola* defence response, like the four genes expressed only in ‘Villard Blanc’. This probably happens because the resistance mechanism is dependent on the activation of R genes, which encode cellular receptors that detect the presence of a particular pathogen, leading to the activation of the signal transduction pathways (Wang et al. 2013). In addition, the susceptibility of this and other species may occur due to failure to assemble the effective defence response, which leads to defective or no recognition of the pathogen (Velasco et al. 2007). Fung et al. (2008) evaluated the interaction between *V. vinifera* and *V. aestivalis* and the *E. necator* pathogen, the agent that causes powdery mildew. These authors observed that for *V. aestivalis* only three genes were regulated as a consequence of the infection, whereas for *V. vinifera*, reprogramming occurred in a high number of genes after pathogen exposure, which is in agreement with the increased expression levels of some genes presented by the ‘Cabernet Sauvignon’ (genes VIT_18s0041g01290, VIT_18s0041g01350, VIT_18s0041g01620, VIT_18s0041g02120 and VIT_18s0041g00090). Another point is that some varieties might present partial resistance to *P. viticola*, which is much more difficult to characterize. A recent work exampled six important components of partial resistance in a diverse set of grapevine varieties leaves showing that partial resistant varieties presented reduced proportion of inoculation, longer latent period, smaller lesion size, fewer production and number of sporangia, shorter infectious period and lower infectivity of sporangia than the susceptible variety ‘Merlot’ (Bove and Rossi 2020). Therefore, evaluating if cultivars and individuals are partially resistant to downy mildew should be a focus of future works.

The early detection of resistance to pathogens in perennial plants is of great importance for breeding programs and for agricultural development. For this objective many studies focus on discovering new and accurate molecular markers to help plant selection. Recently, a study showed the use of haplotype-tagging insertion/deletions (InDels) for downy mildew resistance of grapevine by observing differences in amplicon size between grapes that carry or do not carry Rpv3-1, which can be analyzed by via standard agarose gel electrophoresis or classical melting curve with fluorescent dyes (Foria et al. 2018). In our study, we initially used SSR markers which confirmed that the P3_VVAGL11, VVIN16 and VMC7F2 markers appear to sufficiently co-segregate with the *Rpv3* locus to allow their use as a molecular marker-assisted selection tool for a downy mildew resistance breeding strategy. These three molecular markers have now a dual purpose: they can be used in the previously known diagnosis of the seedlessness character and also in the evaluation of downy mildew resistance in grapevine. A new study also aimed for selecting seedlessness and downy mildew resistance in wild Chinese grapes, by the use of DNA-probe, SSR and SCAR markers (Li et al. 2020). Even though we achieved good results with the SSR, nowadays there are more efficient marker-assisted selection techniques than SSR analysis by polyacrylamide gel. One of these is the KASP ™ genotyping assay. This assay uses a unique form of PCR combined with a homogeneous fluorescence-based information system for the identification and measurement of genetic variation occurring at the nucleotide level to detect single nucleotide polymorphisms (SNPs) or insertions and deletions (INDELs) (He et al. 2014). With the evaluation of our forty-one SNP-based molecular markers, we identified two that are heterozygous and segregate in the ‘Villard Blanc’ self-fertilized population. Markers Rpv3_15 and Rpv3_33 allow for the selection of resistant genotypes by a technique that has several advantages such as a lower genotyping error in positive control DNA samples compared to other techniques and lower per-assay price (7.9-46.1% cheaper) (Semagn et al. 2014). Thus, these markers can generate data on a larger scale, in an automated and cost-efficient fashion (Guimarâes et al. 2009). Moreover, the Rpv3_15 and Rpv3_33 markers hereby presented could be combined with other markers, such as VMC7F2, in marker-assisted breeding programs. Taken together, the results presented in this work contribute with key functional data about the *Rpv3* locus and its associated resistant genes, demonstrating the biotechnological applicability of molecular markers in downy mildew resistance assisted selection strategy for early plant assortment, therefore long term field sustainability.

## Supporting information

STable2

STable3

STable1

## DECLARATIONS

### Availability of data and material

The datasets generated during and/or analyzed during the current study are available in the GenBank repository, https://www.ncbi.nlm.nih.gov/genbank/, with accession number JZ984004 JZ984178, BV005171 and BV140662.

### Author’s contribution statement

L.F.R, J.M., A.W. and V.B. conceived original screenings, research plans, designed experiments and analyzed resulting data; L.F.R and V.B. supervised experiments and writing; J.M., A.W and V.B. performed most experiments; L.F.R and D.D.P provided technical assistance. J.M. and A.W wrote the article with contributions from all authors.

## Ethics declarations

### Ethics approval and consent to participate

Not applicable.

### Consent for publication

Not applicable.

## Supplementary Information (SI)

**Supplementary Fig. 1.**
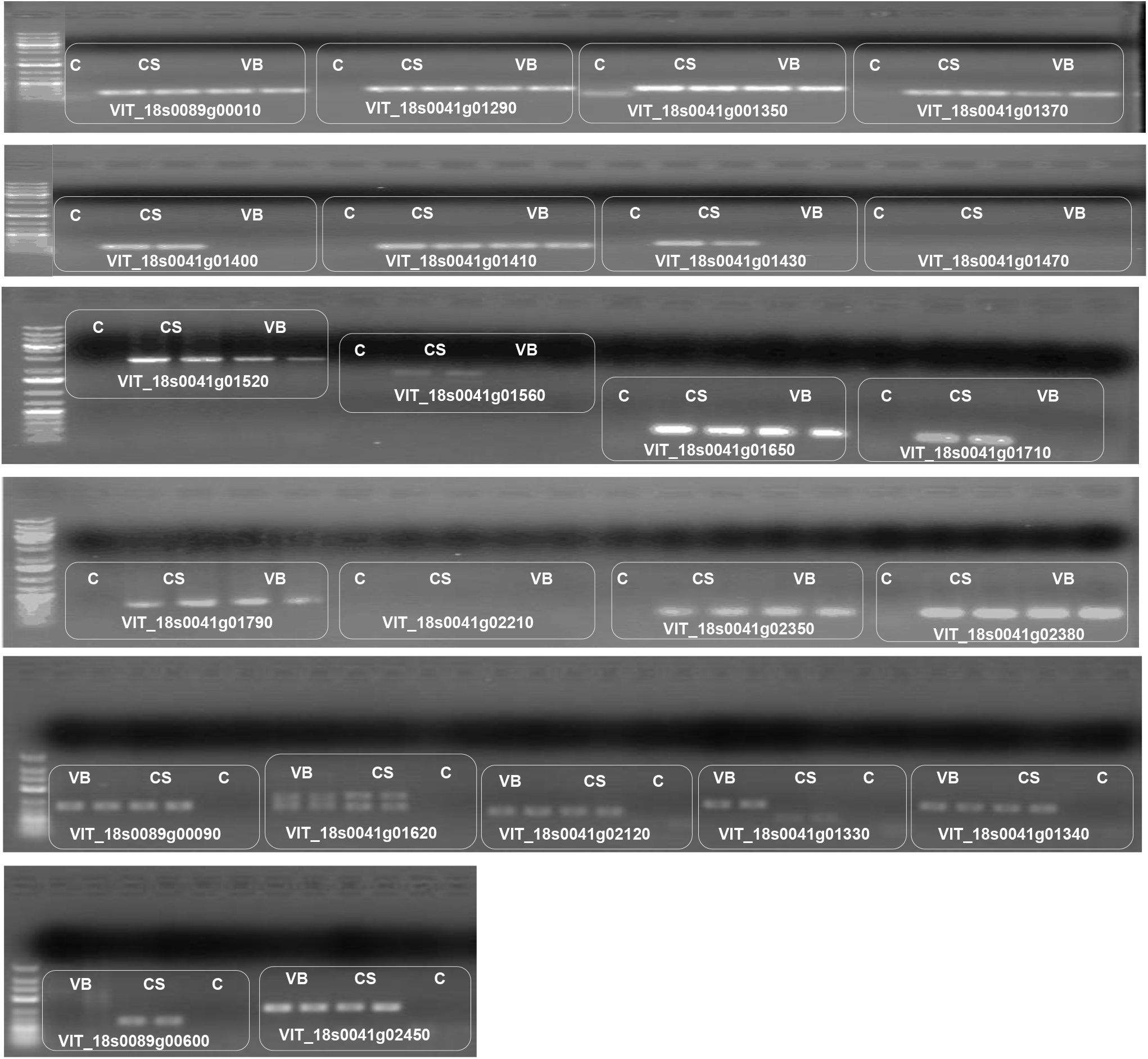
Amplification products of the candidate genes located in *Rpv3* locus. PCR from genomic DNA of the cultivar Villard Blanc (VB) and Cabernet Sauvignon (CS). On the left is the molecular ladder marker (GeneRuler™ Low Range DNA Ladder and 1 Kb Plus DNA Ladder). C = negative control.

**STable.1 Specific primers for candidate genes**

The first column contains the candidate genes names, followed by their specific primers sequence and the expected product size. The Vv04s0044g00580 is the reference gene Actin.

**STable.2 Fisher’s test results**

The first column contains the candidate genes names, followed by their GO terms from the Fisher’s test and by their domain.

**STable.3 Sequences for KASP markers for downy mildew resistance**

## References

Agudelo-Romero P, Erban A, Rego C, et al (2015) Transcriptome and metabolome reprogramming in Vitis vinifera cv. Trincadeira berries upon infection with Botrytis cinerea. J Exp Bot 66:1769–1785. https://doi.org/10.1093/jxb/eru517

Andolfo G, Ercolano MR (2015) Plant innate immunity multicomponent model. Front Plant Sci 6:1–6. https://doi.org/10.3389/fpls.2015.00987

Anonymous (1983) Descriptor list for grapevine varieties and Vitis species 452. Organization Internacional de la Vigne et du Vin (OIV), Paris

Belkhadir Y, Subramaniam R, Dangl JL (2004) Plant disease resistance protein signaling: NBS-LRR proteins and their partners. Curr Opin Plant Biol 7:391–399. https://doi.org/10.1016/j.pbi.2004.05.009

Bellin D, Peressotti E, Merdinoglu D, et al (2009) Resistance to Plasmopara viticola in grapevine “Bianca” is controlled by a major dominant gene causing localised necrosis at the infection site. Theor Appl Genet 120:163–176. https://doi.org/10.1007/s00122-009-1167-2

Bigeard J, Colcombet J, Hirt H (2015) Signaling mechanisms in pattern-triggered immunity (PTI). Mol Plant 8:521–539. https://doi.org/10.1016/j.molp.2014.12.022

Blasi P, Blanc S, Wiedemann-Merdinoglu S, et al (2011) Construction of a reference linkage map of Vitis amurensis and genetic mapping of Rpv8, a locus conferring resistance to grapevine downy mildew. Theor Appl Genet 123:43–53. https://doi.org/10.1007/s00122-011-1565-0

Blum M, Waldner M, Gisi U (2010) A single point mutation in the novel PvCesA3 gene confers resistance to the carboxylic acid amide fungicide mandipropamid in Plasmopara viticola. Fungal Genet Biol 47:499–510. https://doi.org/10.1016/j.fgb.2010.02.009

Bove F, Rossi V (2020) Components of partial resistance to Plasmopara viticola enable complete phenotypic characterization of grapevine varieties. Sci Rep 10:1–12. https://doi.org/10.1038/s41598-020-57482-0

Cabezas JA, Cervera MT, Ruiz-García L, et al (2006) A genetic analysis of seed andberry weight in grapevine. Genome 49:1572–1585. https://doi.org/10.1139/G06-122

Cadle-Davidson L (2008) Variation within and between Vitis spp. for foliar resistance to the downy mildew pathogen Plasmopara viticola. Plant Dis 92:1577–1584. https://doi.org/10.1094/PDIS-92-11-1577

Century KS, Shapiro AD, Repetti PP, et al (1997) NDR1, a pathogen-induced component required for Arabidopsis disease resistance. Science (80-) 278:1963– 1965

Costantini L, Battilana J, Lamaj F, et al (2008) Berry and phenology-related traits in grapevine (Vitis vinifera L.): from Quantitative Trait Loci to underlying genes. BMC Plant Biol 8:1–17. https://doi.org/10.1186/1471-2229-8-38

Costenaro-da-Silva D, Passaia G, Henriques JAP, et al (2010) Identification and expression analysis of genes associated with the early berry development in the seedless grapevine (Vitis vinifera L.) cultivar Sultanine. Plant Sci 179:510–519. https://doi.org/10.1016/j.plantsci.2010.07.021

Creste S, Tulmann Neto A, Figueira A (2001) Detection of single sequence repeat polymorphisms in denaturing polyacrylamide sequencing gels by silver staining. Plant Mol Biol Report 19:299–306. https://doi.org/10.1007/BF02772828

Di Gaspero G, Cipriani G (2003) Nucleotide binding site/leucine-rich repeats, Pto-like and receptor-like kinases related to disease resistance in grapevine. Mol Genet Genomics 269:612–623. https://doi.org/10.1007/s00438-003-0884-5

Di Gaspero G, Copetti D, Coleman C. et al. (2012) Selective sweep at the Rpv3 locus during grapevine breeding or downy mildew resistance. Theor Appl Genet 124, 277–286. https://doi.org/10.1007/s00122-011-1703-8

Díez-Navajas AM, Wiedemann-Merdinoglu S, Greif C, Merdinoglu D (2008) Nonhost versus host resistance to the grapevine downy mildew, Plasmopara viticola, studied at the tissue level. Phytopathology 98:776–780

Doligez A, Bouquet A, Danglot Y, et al (2002) Genetic mapping of grapevine (Vitis vinifera L.) applied to the detection of QTLs for seedlessness and berry weight. Theor Appl Genet 105:780–795. https://doi.org/10.1007/s00122-002-0951-z

Dörmann P, Gopalan S, He SY, Benning C (2000) A gene family in Arabidopsis thaliana with sequence similarity to NDR1 and HIN1. Plant Physiol Biochem 38:789–796

Dry IB, Feechan A, Anderson C, et al (2010) Molecular strategies to enhance the genetic resistance of grapevines to powdery mildew. Aust J Grape Wine Res 16:94–105. https://doi.org/10.1111/j.1755-0238.2009.00076.x

Durrant WE, Dong X (2004) Systemic acquired resistance. Annu Rev Phytopathol 42:185–209. https://doi.org/10.1146/annurev.phyto.42.040803.140421

Eisenmann B, Czemmel S, Ziegler T, et al (2019) Rpv3-1 mediated resistance to grapevine downy mildew is associated with specific host transcriptional responses and the accumulation of stilbenes. BMC Plant Biol 19:1–17. https://doi.org/10.1186/s12870-019-1935-3

Erwin DC, Ribeiro OK (1996) Phytophthora diseases worldwide. American Phytopathological Society (APS Press), St. Paul, Minnesota

Falavigna V da S, Porto DD, Buffon V, et al (2014) Differential transcriptional profiles of dormancy-related genes in ppple buds. Plant Mol Biol Report 32:796–813. https://doi.org/10.1007/s11105-013-0690-0

Figueira D (2013) Cabernet Sauvignon a casta sagrada. In: Prazeres do vinho Fischer BM, Salakhutdinov I, Akkurt M, et al (2004) Quantitative trait locus analysis of fungal disease resistance factors on a molecular map of grapevine. Theor Appl Genet 108:501–515. https://doi.org/10.1007/s00122-003-1445-3

Flor HH (1971) Current status of the gene-fob-gene concept. Annu Rev Phytopathol 3531:275–296

Foria S, Copetti D, Eisenmann B, et al (2020) Gene duplication and transposition of mobile elements drive evolution of the Rpv3 resistance locus in grapevine. Plant J 101:529–542. https://doi.org/10.1111/tpj.14551

Foria S, Magris G, Copetti D, et al (2018) InDel markers for monitoring the introgression of downy mildew resistance from wild relatives into grape varieties. Mol Breed 38:. https://doi.org/10.1007/s11032-018-0880-4

Fröbel S, Zyprian E (2019) Colonization of different grapevine tissues by Pasmopara viticola — a histological study. Front Plant Sci 1–13. https://doi.org/10.3389/fpls.2019.00951

Fung RWM, Gonzalo M, Fekete C, et al (2008) Powdery mildew induces defense-oriented reprogramming of the transcriptome in a susceptible but not in a resistant grapevine. Plant Physiol 146:236–249. https://doi.org/10.1104/pp.107.108712

Garrido, L. da R. Sônego OR (2007) Manejo de doenças da videira. In: Manejo integrado de doenças de fruteiras. Sociedade Brasileira de Fitopatologia, Brasília, pp 65–86

Götz S, García-Gómez JM, Terol J, et al (2008) High-throughput functional annotation and data mining with the Blast2GO suite. Nucleic Acids Res 36:3420–3435. https://doi.org/10.1093/nar/gkn176

Greenberg JT, Vinatzer BA (2003) Identifying type III effectors of plant pathogens and analyzing their interaction with plant cells. Curr Opin Microbiol 6:20–28. https://doi.org/10.1016/S1369-5274(02)00004-8

Grenville-Briggs LJ, Van West P (2005) The biotrophic stages of oomycete-plant interactions. Adv Appl Microbiol 57:217–243. https://doi.org/10.1016/S0065-2164(05)57007-2

Guimarâes CT, Magalhães JV de, Lanza MA, Schyster I (2009) Marcadores moleculares e suas aplicações no melhoramento genético. Inf Agropecuário 30:24–

He C, Holme J, Anthony J (2014) SNP genotyping: the KASP assay. Methods Mol Biol 1145:75–86

Hubank M, Schatz DG (1994) Identifying differences in mRNA expression by representational difference analysis of cDNA. Nucleic Acids Res 22:5640–5648. https://doi.org/10.1093/nar/22.25.5640

IBRAVIN (2012) No Title. In: A Vitivinic. Bras. - Bento Gonçalves INRA, IFV,

SupAgro M (2013) Fiche Variétale Villard Blanc B. Plant Grape

Jaillon O, Aury JM, Noel B, et al (2007) The grapevine genome sequence suggests ancestral hexaploidization in major angiosperm phyla. Nature 449:463–468. https://doi.org/10.1038/nature06148

Jones DA, Takemoto D (2004) Plant innate immunity – direct and indirect recognition of general and specific pathogen-associated molecules. Curr Opin Immunol 16:48– 62. https://doi.org/10.1016/j.coi.2003.11.016

Jones JDG, Dangl JL (2006) The plant immune system. Nature 444:323–239 Kortekamp A (2006) Expression analysis of defence-related genes in grapevine leaves after inoculation with a host and a non-host pathogen. Plant Physiol Biochem 44:58–67. https://doi.org/10.1016/j.plaphy.2006.01.008

Kortekamp A, Zyprian E (2003) Characterization of Plasmopara-Resistance in grapevine using in vitro plants. J Plant Physiol 160:1393–1400. https://doi.org/10.1078/0176-1617-01021

Lefort F, Douglas GC (1999) An efficient micro-method of DNA isolation from mature leaves of four hardwood tree species Acer, Fraxinus, Prunus and Quercus. Ann For Sci 56:259–263

Li S, Liu K, Yu S, et al (2020) The process of embryo abortion of stenospermocarpic grape and it develops into plantlet in vitro using embryo rescue. Plant Cell Tissue Organ Cult 143:389–409. https://doi.org/10.1007/s11240-020-01926-y

Marguerit E, Boury C, Manicki A, et al (2009) Genetic dissection of sex determinism, inflorescence morphology and downy mildew resistance in grapevine. Theor Appl Genet 118:1261–1278. https://doi.org/10.1007/s00122-009-0979-4

Mejía N, Soto B, Guerrero M, et al (2011) Molecular, genetic and transcriptional evidence for a role of VvAGL11 in stenospermocarpic seedlessness in grapevine. BMC Plant Biol 11:1–18. https://doi.org/10.1186/1471-2229-11-57

Merdinoglu D, Wiedeman-Merdinoglu S, Coste P, et al (2003) Genetic analysis of downy mildew resistance derived from Muscadinia rotundifolia. Acta Hortic 603:451–456

Moreira FM, Madini A, Marino R, et al (2011) Genetic linkage maps of two interspecific grape crosses (Vitis spp.) used to localize quantitative trait loci for downy mildew resistance. Tree Genet Genomes 7:153–167. https://doi.org/10.1007/s11295-010-0322-x

Ocarez N, Jiménez N, Núñez R, et al (2020) Unraveling the deep genetic architecture for seedlessness in grapevine and the development and validation of a new set of markers for VviAGL11-based gene-assisted selection. Genes (Basel) 11:1–32

Pfaffl MW (2001) A new mathematical model for relative quantification in real-time RT-PCR. Nucleic Acids Res 29:e45. https://doi.org/10.1093/nar/29.9.e45

Pinto KMS, do Nascimento LC, de Souza Gomes EC, et al (2012) Efficiency of resistance elicitors in the management of grapevine downy mildew Plasmopara viticola: Epidemiological, biochemical and economic aspects. Eur J Plant Pathol 134:745–754. https://doi.org/10.1007/s10658-012-0050-1

Polesani M, Bortesi L, Ferrarini A, et al (2010) General and species-specific transcriptional responses to downy mildew infection in a susceptible (Vitis vinifera) and a resistant (V. Riparia) grapevine species. BMC Genomics 11:1–16. https://doi.org/10.1186/1471-2164-11-117

Possamai T, Migliaro D, Gardiman M, et al (2020) Rpv mediated defense responses in grapevine offspring resistant to Plasmopara viticola. Plants 9:1–10. https://doi.org/10.3390/plants9060781

Reid KE, Olsson N, Schlosser J, et al (2006) An optimized grapevine RNA isolation procedure and statistical determination of reference genes for real-time RT-PCR during berry development. BMC Plant Biol 6:1–11. https://doi.org/10.1186/1471-2229-6-27

Revers LF, Welter LJ, Irala PB, et al (2010) Co-localization of QTLs for seedlessness and downy mildew resistance in grapevine. In: Reisch BI, Londo J (eds) Xth Intl. Conf. on Grapevine Breeding and Genetics. pp 449–456

Ribeiro IJA (2001) Doenças causadas por fungos e bactérias na cultura da videira. In: Boliani AC, CORRÊA L de S (eds) Cultura de uvas de mesa: do plantio à comercialização. ALGRAF, Ilha Solteira, pp 237–239

Ruijter JM, Ramakers C, Hoogaars WMH, et al (2009) Amplification efficiency: linking baseline and bias in the analysis of quantitative PCR data. Nucleic Acids Res 37:1–12. https://doi.org/10.1093/nar/gkp045

Saifert L, Sánchez-Mora FD, Assumpção WT, et al (2018) Marker-assisted pyramiding of resistance loci to grape downy mildew. Pesqui Agropecu Bras 53:602–610. https://doi.org/10.1590/S0100-204X2018000500009

Salmaso M, Malacarne G, Troggio M, et al (2008) A grapevine (Vitis vinifera L.) genetic map integrating the position of 139 expressed genes. Theor Appl Genet 116:. https://doi.org/10.1007/s00122-008-0741-3

Sargolzaei M, Maddalena G, Bitsadze N, et al (2020) Rpv29, Rpv30 and Rpv31: Three Novel Genomic Loci Associated With Resistance to Plasmopara viticola in Vitis vinifera. Front Plant Sci 11:1–16. https://doi.org/10.3389/fpls.2020.562432

Schwander F, Eibach R, Fechter I, et al (2012) Rpv10: A new locus from the Asian Vitis gene pool for pyramiding downy mildew resistance loci in grapevine. Theor Appl Genet 124:163–176. https://doi.org/10.1007/s00122-011-1695-4

Semagn K, Babu R, Hearne S, Olsen M (2014) Single nucleotide polymorphism genotyping using Kompetitive Allele Specific PCR (KASP): overview of the technology and its application in crop improvement. Mol Breed 33:1–14. https://doi.org/10.1007/s11032-013-9917-x

Soanes DM, Talbot NJ (2008) Moving targets: rapid evolution of oomycete effectors. Trends Microbiol 16:507–510. https://doi.org/10.1016/j.tim.2008.08.002

Takken FL, Albrecht M, Tameling W IL (2006) Resistance proteins: molecular switches of plant defence. Curr Opin Plant Biol 9:383–390. https://doi.org/10.1016/j.pbi.2006.05.009

Tessmann DJ, Vida JB (2005) Principais doenças fúngicas da videira. Rev Atualidades Agrícolas 14–15

Toffolatti SL, Maddalena G, Maghradze D, Bianco PA (2016) Evidence of resistance to the downy mildew agent Plasmopara viticola in the Georgian Vitis vinifera germplasm. https://doi.org/10.5073/vitis.2016.55.121-128

Unger S, Büche C, Boso S, Kassemeyer HH (2007) The course of colonization of two different Vitis genotypes by Plasmopara viticola indicates compatible and incompatible host-pathogen interactions. Phytopathology 97:780–786. https://doi.org/10.1094/PHYTO-97-7-0780

Velasco R, Zharkikh A, Troggio M, et al (2007) A high quality draft consensus sequence of the genome of a heterozygous grapevine variety. PLoS One 1–18. https://doi.org/10.1371/journal.pone.0001326

Venuti S, Copetti D, Foria S, et al (2013) Historical introgression of the downy mildew resistance gene Rpv12 from the Asian species Vitis amurensis into grapevine varieties. PLoS One 8:e61228. https://doi.org/10.1371/journal.pone.0061228

Wang P, Liu C, Wang D, et al (2013) Isolation of resistance gene analogs from grapevine resistant to downy mildew. Sci Hortic (Amsterdam) 150:326–333. https://doi.org/10.1016/j.scienta.2012.11.035

Welter LJ, Göktürk-Baydar N, Akkurt M, et al (2007) Genetic mapping and localization of quantitative trait loci affecting fungal disease resistance and leaf morphology in grapevine (Vitis vinifera L). Mol Breed 20:359–374. https://doi.org/10.1007/s11032-007-9097-7

Winter D, Vinegar B, Nahal H, et al (2007) An “electronic fluorescent pictograph” Browser for exploring and analyzing large-scale biological data sets. PLoS One 2:. https://doi.org/10.1371/journal.pone.0000718

Zeng Y, Yang T (2002) RNA isolation from highly viscous samples rich in polyphenols and polysaccharides. Plant Mol Biol Report 20:417a–417e. https://doi.org/10.1007/BF02772130

Zyprian, E, Ochßner I, Schwander F. et al. (2016) Quantitative trait loci affecting pathogen resistance and ripening of grapevines. Mol Genet Genomics 291, 1573– m1594. https://doi.org/10.1007/s00438-016-1200-5

